# A FIJI Macro for quantifying pattern in extracellular matrix

**DOI:** 10.1101/867507

**Authors:** Esther Wershof, Danielle Park, David J Barry, Robert P Jenkins, Antonio Rullan, Anna Wilkins, Ioannis Roxanis, Kurt I Anderson, Paul A Bates, Erik Sahai

## Abstract

Diverse extracellular matrix patterns are observed in both normal and pathological tissue. However, most current tools for quantitative analysis focus on a single aspect of matrix patterning. Thus, an automated pipeline that simultaneously quantifies a broad range of metrics and enables a comprehensive description of varied matrix patterns is needed. To this end we have developed an ImageJ plugin called TWOMBLI, which stands for <u>T</u>he <u>W</u>orkflow <u>O</u>f <u>M</u>atrix <u>B</u>io<u>L</u>ogy <u>I</u>nformatics. TWOMBLI is designed to be quick, versatile and easy-to-use particularly for non-computational scientists. TWOMBLI can be downloaded from https://github.com/wershofe/TWOMBLI together with detailed documentation. Here we present an overview of the pipeline together with examples from a wide range of contexts where matrix patterns are generated.

## Introduction

The extracellular matrix (ECM) provides support and structure to multicellular organisms and also guides the migration of cells ^1,2^, including leukocytes engaged in immune surveillance ^3^. Changes in the ECM are central to the aging process, the ECM is re-built and remodelled in response to tissue damage, and it is further altered in pathologies such as cancer. The architecture of the ECM has lately been the subject of renewed focus ^4–6^, however standardised quantification of ECM patterns, particularly in pathological decision-making is lacking. The generation of metrics that describe ECM pattern could lead to insights into a wide range of fields, ranging from experimentalists interested in cell migration and remodelling of matrices to clinicians researching conditions such as cancer and fibrosis.

Diverse extracellular matrix patterns are observed in both normal and pathological tissue. Numerous metrics have already been applied to such ECM images. These range from simple abundance and area measurements to more complex textural features such as grey level co-occurrence matrices^7^ and fractal dimension^8^. One commonly used metric is matrix fibre alignment, which is known to be a promoter of cancer cell invasion ^9–11^. A pipeline already exists in MATLAB for quantifying alignment of ECM ^12^, and a further extension of this work enables alignment to be related to the tumour margin^13^. OrientationJ is an automated ImageJ plugin that is able to create vector fields and perform directional analysis on fibres, but this does not lend itself to quantifying overall matrix patterns. A number of studies have attempted to quantify additional matrix metrics, but these typically require heavy manual intervention ^14,15^. Furthermore, each tool typically only generates metrics relating to one aspects of ECM organisation. This makes collating a broad range of metrics for analysis both challenging and time-consuming. There is a need for an end-to-end pipeline for quantifying ECM patterns, which is automated and easy-to-use on versatile data sets. To this end, we have created the ImageJ macro plugin TWOMBLI, which stands for <u>T</u>he <u>Wo</u>rkflow <u>O</u>f <u>M</u>atrix <u>B</u>io<u>L</u>ogy Informatics. The aim of TWOMBLI is to quantify matrix patterns in an ECM image by deriving a range of metrics, which can then be analysed in conjunction with clinical data if desired.

## Results

We sought to generate a tool for analysis of ECM patterns, whereby a user could enter varied matrix images into the pipeline and derive a meaningful quantification of a variety of matrix features as output (Figure 1). In particular, we were keen that a broad range of metrics would capture diverse features of ECM patterns. FIJI was employed as the supporting platform for the plugin, since it enabled us to build on many existing tools and downloads such as Ridge Detection^16^, Anamorf^17^, OrientationJ, and BIOP and is also familiar to many scientists and clinicians.

**Figure 1:**
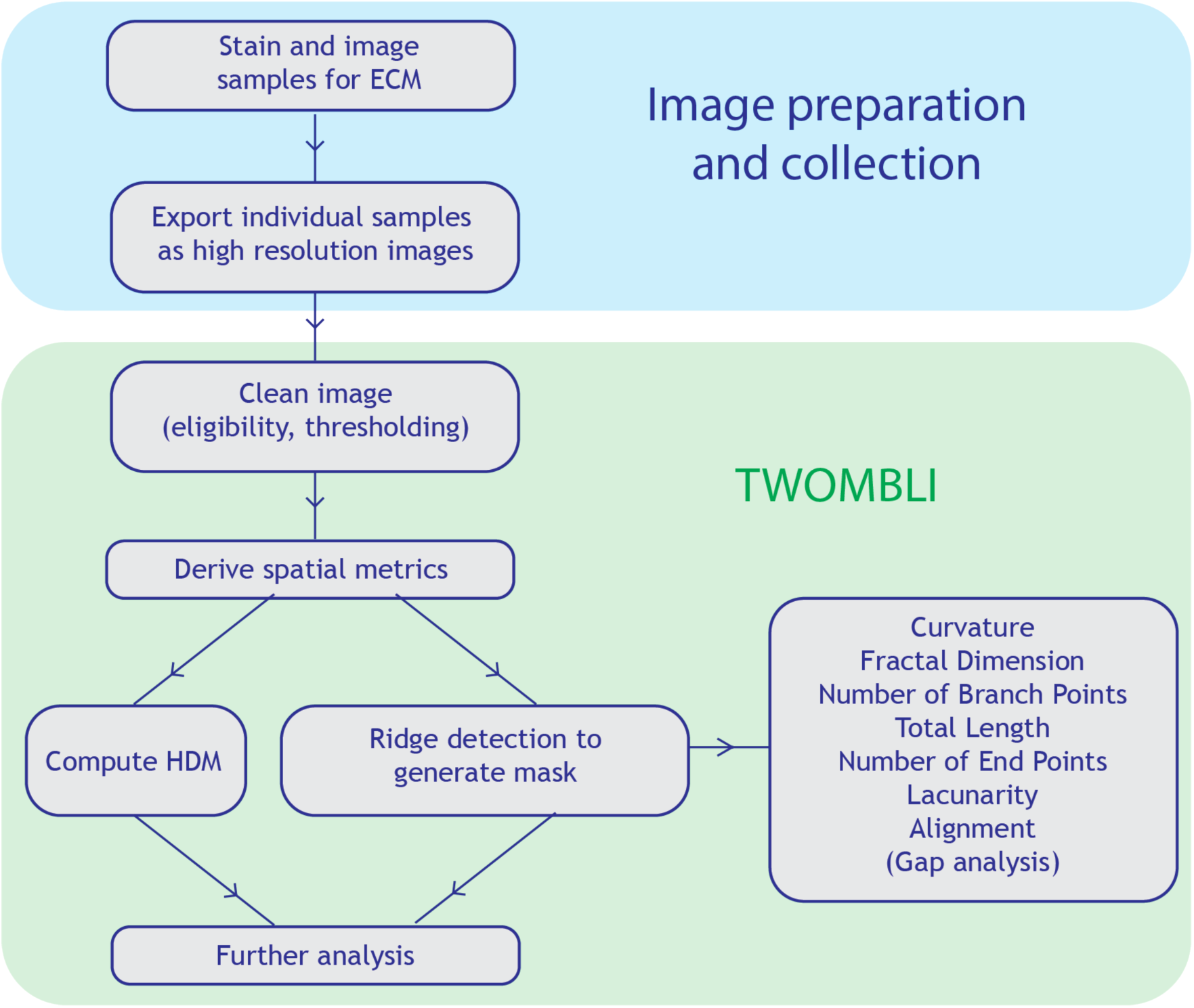
Workflow diagram of quantification of matrix patterns. End-to-end pipeline from obtaining the samples through matrix quantification to survival analysis based on this matrix metrology. A list of metrics is given in the right-most box.

### Development of the TWOMBLI pipeline

TWOMBLI exists as a user-friendly ImageJ plugin that can be downloaded with detailed documentation from https://github.com/wershofe/TWOMBLI. TWOMBLI relies heavily on existing ImageJ plugins Ridge Detection ^16^ and AnaMorf ^17^, providing an end-to-end pipeline for users. The user can input tissue samples stained for ECM components and is then guided through image pre-processing before finally being given a comprehensive output of ECM metrics in a single csv file with accompanying processed images. Central to this process is the generation of mask files of the matrix network using the Ridge Detection tool. Ridges and ravines in the image are detected from the spatial derivatives of the image. The algorithm carries out simple local searches of the spatial derivative information in order to construct lines, making the algorithm far more computationally efficient than alternative approaches. The algorithm also allows for subpixel detection of lines and is robust to an asymmetric contrast gradient on either lateral edge of the line allowing for good line extraction even in low resolution images. The mask files are generated as outputs of the pipeline to enable visual verification of correct thresholding. Furthermore, an additional script is included in the TWOMBLI repository for optional cropping down of images to regions of interest. The principal stages of TWOMBLI are: Prechecking, pre-processing and processing. The prechecks (steps 0-3) are carried out manually by the user to check the eligibility of potential input images following prompts. An image needs to be in focus, have high enough resolution and not contain too many artefacts or regions that are not pertinent for the analysis. In pre-processing (steps 4-10), the user is guided through selecting a subset of test images and then choosing appropriate parameters for thresholding these test images. In addition to contrast saturation, which relates obtaining the appropriate contrast between areas of matrix and no matrix for subsequent steps, a line width parameter is requested. This value sets the thickness of matrix fibres that will be identified. These parameters are saved so that in future runs, the user can skip directly to the processing stage. In processing (steps 11-14), all of the images are analysed in batch using the parameters generated in the pre-processing stage. An input image of 1MB in size takes approximately 30 seconds to process on a computer with 2.2 GHz Intel Core i7 and 16GB of memory. For tips on handling larger files and other troubleshooting see Documentation.

### Metrics for matrix quantification

We selected that covered different aspects of ECM pattern (a schematic of some key metrics is shown in Figure 2). These metrics can be split between those describing individual fibres and those describing general ECM pattern: number of fibre end points, number of fibre branchpoints, total length of fibres, fibre curvature, alignment, proportion of high-density matrix, ECM fractal dimension, hyphal growth unit (a measure of the number of end points per unit length), matrix gaps, and lacunarity (a measure of how the ECM fills the space) ^17^. These metrics together could characterise different matrix properties. Apart from high density matrix, all metrics were calculated from the mask images generated within the TWOMBLI pipeline.

**Figure 2:**
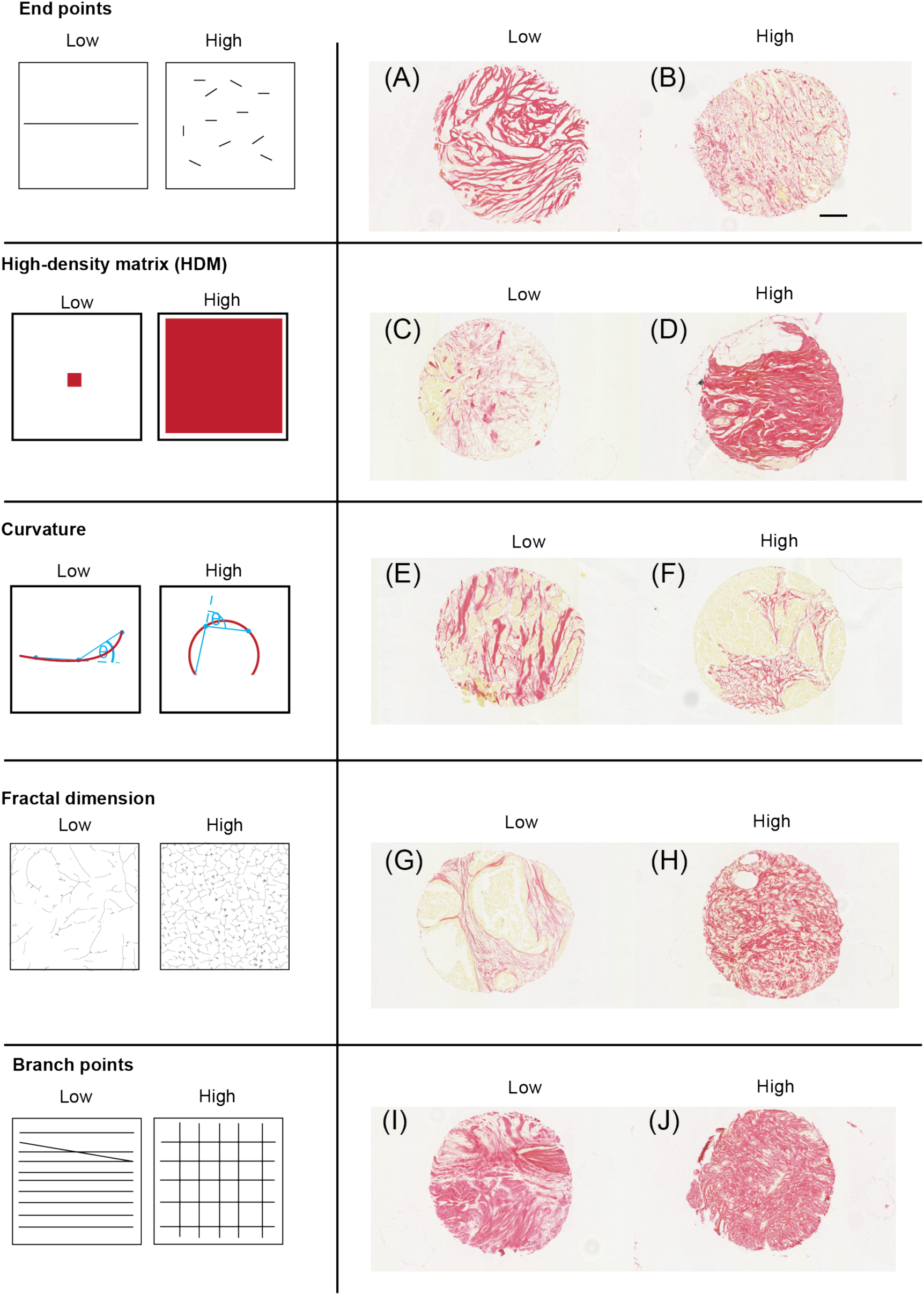
Schematics and example tissue biopsies of ECM metrics. (A)-(J) Images show picrosirius red staining breast cancer biopsies. Each biopsy is 600 microns in diameter, scale bar is 100 microns.

#### Quantification of individual fibres

Number of endpoints is an intuitive count of the number of ends of the fibres or filaments in the mask image. Number of branchpoints is the number of intersections of mask fibres in the image. Total length is the sum of the length of all mask fibres in the image. This quantity can be useful in normalising the number of branchpoints and endpoints. The hyphal growth unit (HGU) corresponds to the number of end points per unit length. Curvature was measured as the mean change in angle moving incrementally along individual mask fibres by user-specified windows.

#### Quantification of global pattern

High-density matrix (HDM) is a measure of the proportion of pixels in an image corresponding to matrix, as defined by the user-specified contrast saturation parameter and subsequent thresholding. The alignment metric captures the extent to which fibres within the field of view are oriented in a similar direction^15,18^. It is calculated from the global gradient structure tensor defined as

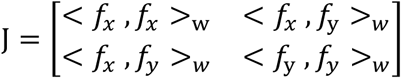

where *f*_*x*_ and *f*_*y*_ are partial spatial derivatives of the image, f(x, y), and

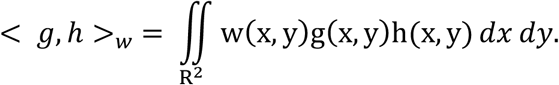

The function, *w*(*x, y*) is a normalized weighting window centred on the region of interest. Alignment is then determined from the coherency metric

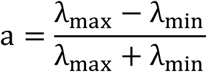

where λ_min_ is the smallest eigenvalue (minor axis) and λ_max_ is the largest eigenvalue (major axis) of the tensor, yielding a value in the range [0,1], where zero represents complete isotropy and one represents perfect alignment. Fractal dimension is an indicator of the self-similarity and complexity of the ECM and is bound between in the range [1,2] for a single 2D image slice. Specifically, the metric used is the box-counting dimension ^19^. A grid with squares of side length *ϵ* is overlaid over the image. The number of squares *N*(*ϵ*) which are occupied by the non-background part of the image N is recorded. As *ϵ* gets smaller, *N*(*ϵ*) increases. Fractal dimension is then computed as the limit of the following equation:

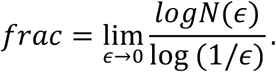

The lacunarity metric reflects number and size of gaps in the matrix. Briefly, the variation in pixel intensity is sampled in different directions and in different size grids and a single average value is returned, with larger values indicated larger space in the matrix pattern. Formally, lacunarity is quantified as

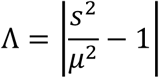

Where *μ* and *s* are respectively the mean and standard deviation of grey level enclosed within a region ^20^. In addition, an optional step in the plugin allows the user to perform gap analysis on matrix patterns using the Max Inscribed Circle function available from the BIOP plugin in FIJI. The algorithm analyses spaces between objects (in this case, the fibres in the masks derived by TWOMBLI) by fitting circles of decreasing radius to fill in the gaps ^21^. This information is reported in individual .csv files containing the size of all gaps identified, which allows for individual researchers to choose whether to focus the average size of gaps, the shape of the distribution, or even the size of gaps in the tails of the distribution. Depending on the tissue and context, these different metrics of gap sizes provide additional insight into the structure of the matrix patterns.

### Use of the TWOMBLI pipeline

Figure 3A shows an example of user input together with the different outputs from the TWOMBLI pipeline. The example shown involves Picrosirius Red staining of fibrillar collagen in a breast cancer biopsy, with the matrix filaments identified shown in the right-hand image. Importantly, the pipeline can be used to analyse images of ECM acquired via a wide variety of imaging techniques and of varying file types. Figure 3B shows its application to cell-derived matrix assays (both isotropic and anisotropic), second-harmonic imaging of mouse tissues, and synthetic matrix patterns generated via computer simulations ^5^. In all examples, the algorithm is able to correctly identify matrix filaments.

**Figure 3:**
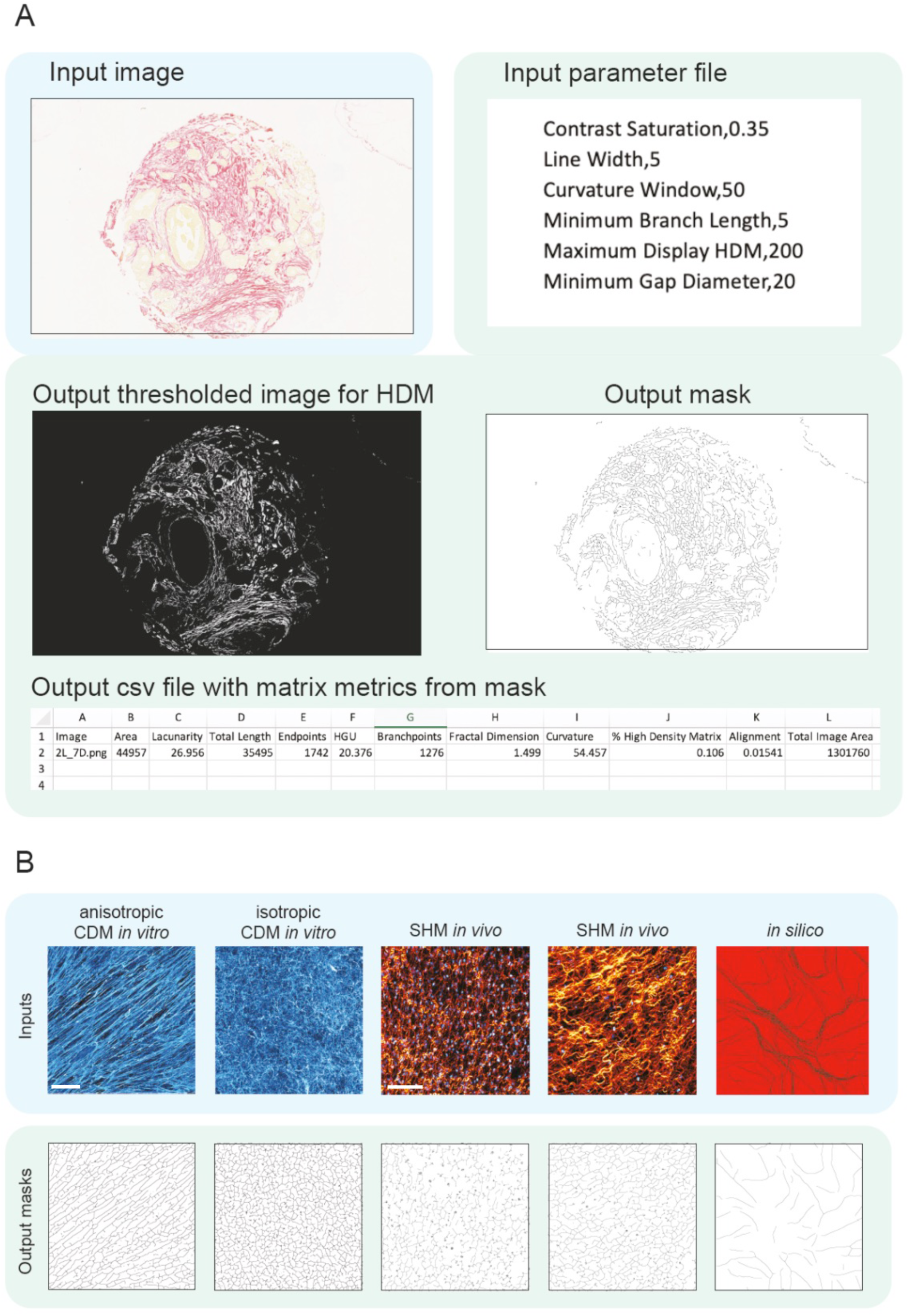
Input and outputs of TWOMBLI pipeline. (A) User inputs an image of a sample stained for ECM components (in this case, collagen is stained with picrosirius red). Outputs consist of a mask, a csv file containing matrix metrics based on the mask and a processed image thresholded for HDM. (B) Example input images can be acquired from a range of sources: for example (from left to right), fibronectin-stained CDM in vitro (images are 500 microns x 500 microns, fibronectin in blue), second harmonic imaging in vivo (images are 400 microns x 400 microns, collagen in orange and fibroblast nuclei in blue), and in silico ECM. Scale bars are 100 microns.

The utility of quantification tools is crucially dependent upon their robustness to variations in the exact details of data acquisition. To explore this, we captured images of regions of two cell-derived matrices with different features: one was largely isotropic with finer fibres (hereafter referred to as isotropic, Supplementary Figure 1A) and the other more anisotropic with thicker, sparser, fibres (hereafter referred to as anisotropic, Supplementary Figure 1A). Images of these two matrices were acquired using different microscope objectives, gain settings, pixel sizes, and focal planes (different images were generated from either single confocal sections of a CDM taken a different focal planes, a single confocal section with a wide pinhole setting, or a z projection of a stack of confocal sections). These images were then analysed using two different line width settings in TWOMBLI. In addition, we electronically degraded the quality of some images using Gaussian blur functions of either 2 or 3 pixels radius or the standard noise function in ImageJ. This set of images were then run through TWOMBLI. Table S1 shows the output metrics and Supplementary Figure 1B shows a subset of the mask images for the more isotropic matrix. The absolute metrics for filament length, endpoints and branchpoints scaled with the size of the input image in pixels – this is also clearly visible in the masks that are generated, with reduced matrix network complexity apparent as the number of pixels is reduced. Normalisation of the filament length to the image size and the endpoints and branchpoints to the total filament length eliminated the relationship between image size (in pixels) and length, endpoints or branchpoints (Table S1 and Supplementary Figure 2). It should be noted that the normalised endpoint metric is the inverse of the HGU metric. Therefore, we use normalised versions of these metrics in subsequent analyses and would recommend that other users do the same.

Having determined which metrics required normalisation and which did not, we then asked whether TWOMBLI could reliably quantify differences between the isotropic and anisotropic CDMs. To obtain a broad overview of the data, we generated a PCA plot using the normalised and scale-free metrics, with the exception of curvature that exhibited high variability. Figure 4A shows that the quantification of the isotropic CDM and anisotropic CDM largely fall into two distinct clusters; however, there are some outliers and the separation is not perfect. Further inspection revealed that the most marked outliers were under-exposed images or those with added Gaussian blur or noise (Table S1). If the analysis was restricted to images captured using the same 20x objective, with appropriate exposure, the same pixel size, and same line width value, then TWOMBLI generated two well-segregated clusters in PCA analysis corresponding to the isotropic and anisotropic matrices (Figure 4B). Analysis of individual variables revealed that many were remarkably robust to the image acquisition settings, with alignment, fractal dimension, and lacunarity being particularly insensitive to variation in microscope settings and even the use of a 10x or 20x objective (Figure 4C and Table S1). Curvature was the one parameter that exhibited a high degree of variation. Closer inspection of the details of pixel size and line width selected revealed that the normalised endpoint metric was sensitive to the choice of line width, with more nuanced variation in other metrics depending on the pixel size and line width choice. For optimal robustness, we would recommend that users select a single line width for their analysis and do not vary this. Overall, these data indicate that the metrics generated TWOMBLI are robust to the precise choice of microscope objective, pixel size, and line width. The biggest factor leading to variation in output metrics was incorrect image exposure. The precise choice of focal plane for imaging and its thickness had relatively little effect. Minimal variation in metrics was achieved with a consistent image resolution and line width choice.

**Figure 4:**
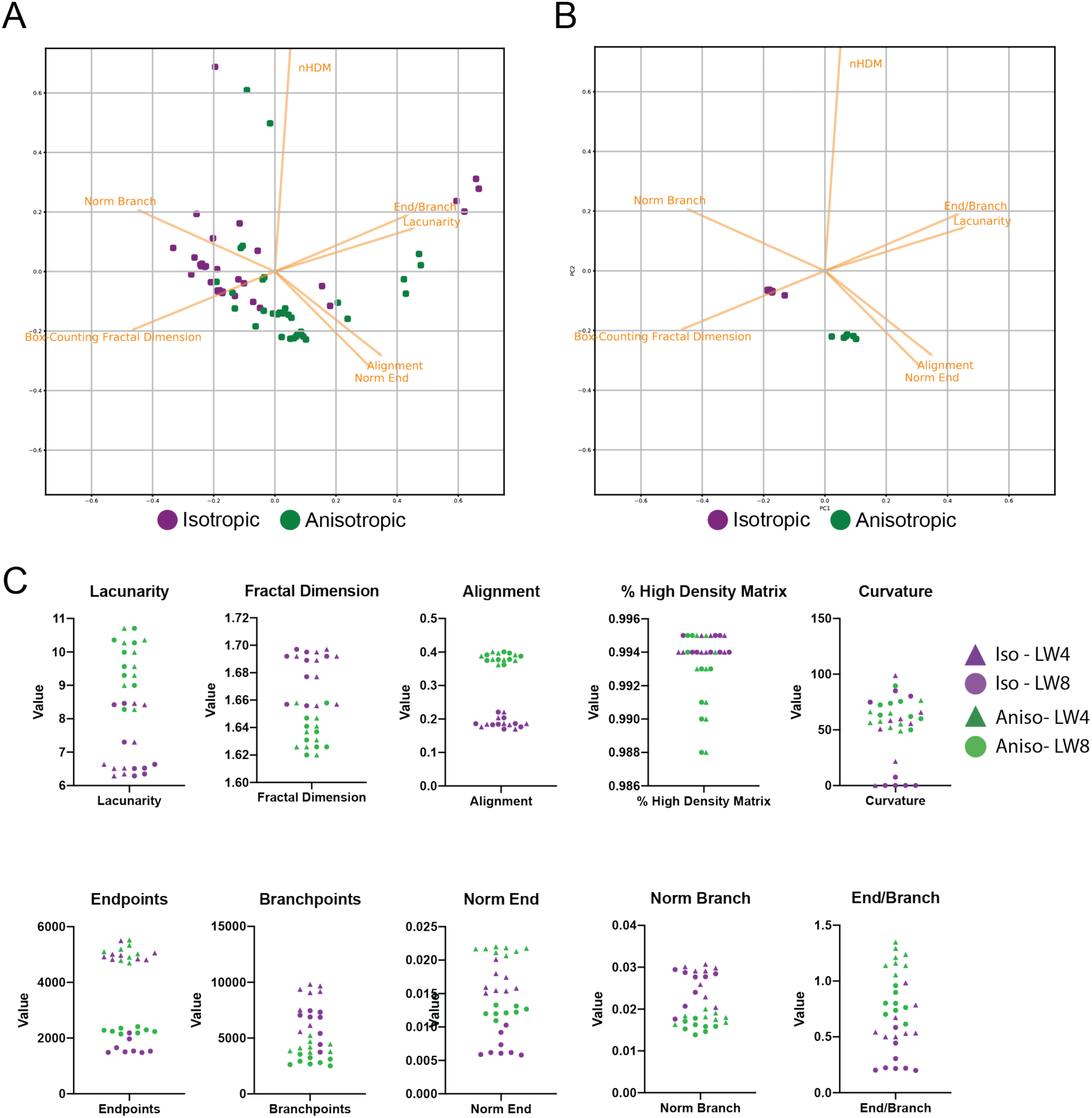
Robustness of matrix metrics. (A) Image shows a PCA plot of both experimental and artificial variations in images of the same region of isotropic cell-derived matrix (purple dots) and anisotropic cell-derived matrix (green dots). (B) Image shows a PCA plot of metrics derived from images captured with a 20x objective of uniform pixel size, exposure, and line width of the same region of isotropic cell-derived matrix (purple dots) and anisotropic cell-derived matrix (green dots). (C) Plots show the values of the indicated metrics with isotropic CDM indicated in purple and anisotropic CDM labelled in green. As in (B) only images with the same pixel size and appropriate exposure are included, but images captured with a 10x objective and more divergent focal planes are also included. Furthermore, two different line width settings are shown: triangles denote use of line width 4 and circles use of line width 8.

### Output metrics are able to distinguish between different matrix patterns

Having determined that TWOMBLI is able to generate reliable metrics, so long as simple principles of consistency in image acquisition and analysis parameters were followed, we tested whether TWOMBLI could provide a quantitative framework for discriminating between different ECM architectures. To this end, images of cell-derived matrices (CDM) generated by seven different isolates of fibroblasts were analysed, with multiple images for each example. Figure 5A shows that TWOMBLI could effectively identify matrix fibres. Furthermore, the alignment metric accurately separated the CDMs which had previously been classified as either isotropic and anisotropic using a more complex MATLAB tool^12^. TWOMBLI also revealed notable differences in addition to alignment, the larger gaps in the MAF2 matrix were reflected in the lower fractal dimension and higher lacunarity (Figure 5B and Table S2). As might be expected, isotropic matrix patterns generally had lower levels of normalised branchpoints. PCA analysis of the different CDMs is presented in Supplementary Figure 3.

**Figure 5:**
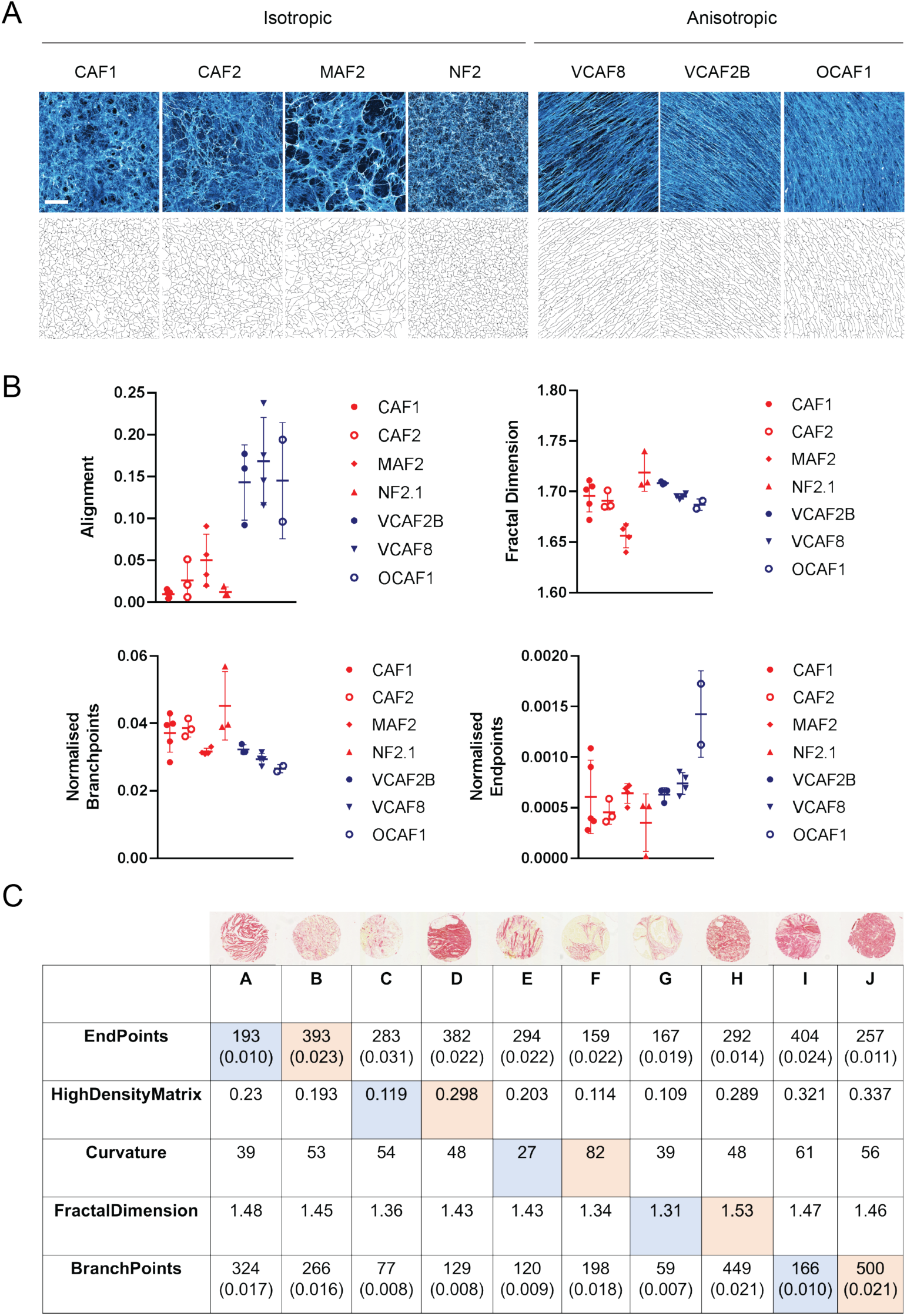
Using TWOMBLI for quantification of ECM patterns (A) CDM imaging of fibronectin produced by seven different fibroblast lines in vitro ^4^. Four patterns are isotropic (left) and three patterns are anisotropic (right). Images are 500 microns x 500 microns, scale bar is 100 microns. Corresponding masks derived in TWOMBLI are displayed underneath. (B) Quantification of matrix patterns from (A) across a number of metrics. Normalisation is performed by dividing the raw value by the total length of fibres in the masks. (C) Output metrics for the corresponding tumour biopsies shown in Figure 2. Pairs (A, B), (C, D) etc. show contrasting biopsies with low and high values of each metric as indicated in the table in blue/orange respectively. For endpoints and branchpoints, normalised values are given in brackets below the raw value. The normalised value is computed by dividing the raw value by the total length of the fibres in the mask.

Quantification of matrix from the breast cancer biopsies presented in Figure 2 shows how some metrics such as high-density matrix (HDM) are intuitively different, whilst other metrics such as curvature and branchpoints are more nuanced (Figure 5C). This suggests that, in addition to quantifying readily discernible pattern features, TWOMBLI could be particularly helpful in identifying patterning properties that are difficult to ascertain by eye. The application of TWOMBLI to prostate biopsies is demonstrated in Supplementary Figure 4A, with quantitative differences in high density matrix, fractal dimension, and lacunarity collectively distinguishing tumour regions from both normal glandular and normal stromal regions of prostate tissue (Supplementary Figure 4B and Table S3). Taken together with the robustness results, these analyses suggest that TWOMBLI is able to distinguish between clearly different matrix patterns.

Finally, we sought to test whether our pipeline might have utility beyond the analysis of extracellular matrix fibres. Dynamic filamentous polymers, including F-actin and microtubules, within cells collectively make up the intracellular cytoskeleton. Supplementary Figure 5 shows images of the F-actin network within the NF2 and CAF1 fibroblasts used to generate the CDMs that we analysed in Figure 5. We have previously reported increased F-actin stress fibres in CAF1 and, consistent with this, the TWOMBLI pipeline correctly identifies an increased length of filamentous structures. This demonstrates that our analysis tool could analyse diverse imaging data spanning sub-cellular filamentous networks through to the extracellular matrix in pathological samples.

## Discussion

Much work has focused on identifying patterns of cells in tissues ^22–24^. However, ECM organisation, which plays a crucial part in tissue architecture, has been somewhat neglected. We have developed TWOMBLI to quantify a wide range of matrix features, enabling further understanding into the relevance of ECM organisation in a wide range of contexts, from tissue damage ^25–27^ to ageing ^28^, development ^29^ to fibrotic disease and cancer. This tool is designed to be a ‘one stop shop’ that generates a wide range of metrics that capture diverse aspects of matrix pattern in the same pipeline, thereby saving researchers from having to run multiple analytical tools in parallel on the same images. FIJI was chosen as the platform because it is freely available, well known and widely used by the biological and medical sciences community ^4,30^

Our analysis indicates that the metrics generated are relatively robust to both the precise region of matrix selected and to the exact microscope settings (Figure 4). Sub-optimal exposure of images was the most detrimental factor in terms of the robustness of the output metrics generated and we would advise researchers to ensure appropriate exposure settings. The objective used and pixel size should be kept constant for optimal results. Slight variations in focal plane or the thickness of the optical section had only minor effects on the output metrics (Figure 4B). Attention should be paid to the ridge detection width because this parameter dictates whether the algorithm identifies fine or thick matrix fibres. Analysis of the same images using two different line widths revealed that the identification of endpoints and branchpoints was sensitive to the choice of line width. However, many parameters were relatively insensitive to this setting, which should enable comparison of metrics between different studies conducted using slightly different analysis settings. The only parameter that we would advise extra caution to be taken over is curvature, which showed a high degree of variation. In some cases, it may be advantageous to run the same images with two different ridge detection widths so that both fine and coarse features can be captured.

The tool that we describe here is designed to be simple to use and generate a wide range of metrics. It is not designed to be highly specialised in its handling of particular matrix features. Excellent specialist tools exist for researchers who may wish to dig deeper into particular features of the ECM; in particular, the Eliceiri group has developed a suite of tools to interrogate matrix alignment using MATLAB^13^, including analysis of matrix fibre orientation relative to tumour cells. Our group has also generated more bespoke tools for measuring matrix alignment over a range of length scales and this may be of use to researchers wishing to compare short-range and long-range matrix alignment ^4,5^. The alignment metric in TWOMBLI is not capable of this level of sophistication and simply generates a global alignment score for the whole field of view. MATLAB tools for the analysis of matrix gaps have also been generated ^30^. Haralick features have also been used to measure ECM organisation^7^; however, we did not include it because textural analysis is not well-suited to our filament-based approach. Nonetheless, Haralick analysis may be a useful addition to future iterations of matrix analysis platforms, especially if performed on images prior to filament tracing.

The tool that we describe has been developed with the purpose of quantifying ECM fibres, it can also be applied to other classes of biological images, such as networks of F-actin or microtubules or even vascular networks. Its broad applicability is evidenced by its ability to analyse the F-actin cytoskeleton of the fibroblasts (Supplementary Figure 5). Furthermore, the ability of this tool to segregate normal prostate images from tumour regions suggests that it could be exploited in the analysis of clinical samples. It is our sincere hope that this versatile tool will be of use to cell biologists, tissue biologists, and pathologists, enabling these communities to study the consequences of different matrix architectures in disease.

## Methods

### Computational analysis

All computational methods are described in detail in the TWOMBLI documentation which can be found at https://github.com/wershofe/TWOMBLI.

### Picrosirius red staining

Samples were stained using ABCAM ab150681 Picrosirius Red Kit. Briefly, slides were deparaffinised and hydrated, before applying Picro-Sirius Red Solution for 60 minutes, rinsing twice in acetic acid, then alcohol dehydration, and finally mounting. Slides were then scanned at 10x using Zeiss Axio Scan.Z1.

### Cell-derived matrix assay

The CDM assay and the fibroblasts used are described in Park et al ^4^. Briefly, glass-bottom dishes (MatTek, P35-1.5-14-C) were pre-prepared with 0.2% gelatin solution (1h, 37 °C), then 1% glutaraldehyde for 30 min at room temperature (RT). After PBS buffer solution wash, the plate was incubated with 1 M ethanolamine for 30 min (RT). After two washes with PBS we seeded 7 × 104 cells in media with 100 μg ml−1 ascorbic acid ((+)-sodium L-ascorbate, Sigma, A4034). Cells were kept for 6 days and media changed every 2 days. We used extraction buffer and washed several times with PBS before immunofluorescence for ECM using anti-fibronectin-FITC (1:50 dilution; Abcam, ab72686). Samples were imaged using a Zeiss LSM 780 microscope using either a 20x 0.75NA objective or 10x 0.45NA objective.

### Collagen Imaging

Second harmonic generation imaging for collagen was performed as described in Park et al using a Zeiss LSM 780 microscope with Mai Tai multi-photon laser. Fresh post-mortem tissue from PDGFRA::H2B-eGFP mice was used.

## Supplementary Information

Supplementary Text 1 = TWOMBLI Documentation

Supplementary Video = TWOMBLI Tutorial

## Acknowledgements

We thank members of the Sahai group, Bella Kotantaki, Florian Laflorêts for advice and comments. We thank Prof. David Dearnley, Prof. Emma Hall, Dr Michael Toss, and Dr Emad Rakha for assistance with clinical samples. E.W., D.P., D.B., A.W., R.P.J., K.I.A., P.A.B., and E.S. were funded by the Francis Crick Institute, which receives its core funding from Cancer Research UK (FC001003, FC001144), the UK Medical Research Council (FC001003, FC001144) and the Wellcome Trust (FC001003, FC001144). A.R. was supported by the Spanish Society for Medical Oncology (Beca Fundación SEOM). E.S and D.P. also received funding from Breast Cancer Now (2013NovPR182). A.W. is also supported by a Crick i2i grant.

## Author contributions

E.W., D.P., P.A.B., and E.S. conceived the study. E.W. wrote the code, with assistance from D.B. and advice from R.P.J. A.W., A.R., and I.R. enabled access to and analysed the patient samples. E.W. and E.S. wrote the manuscript with editorial assistance from all authors.

---

**Supplementary Figure 1.**
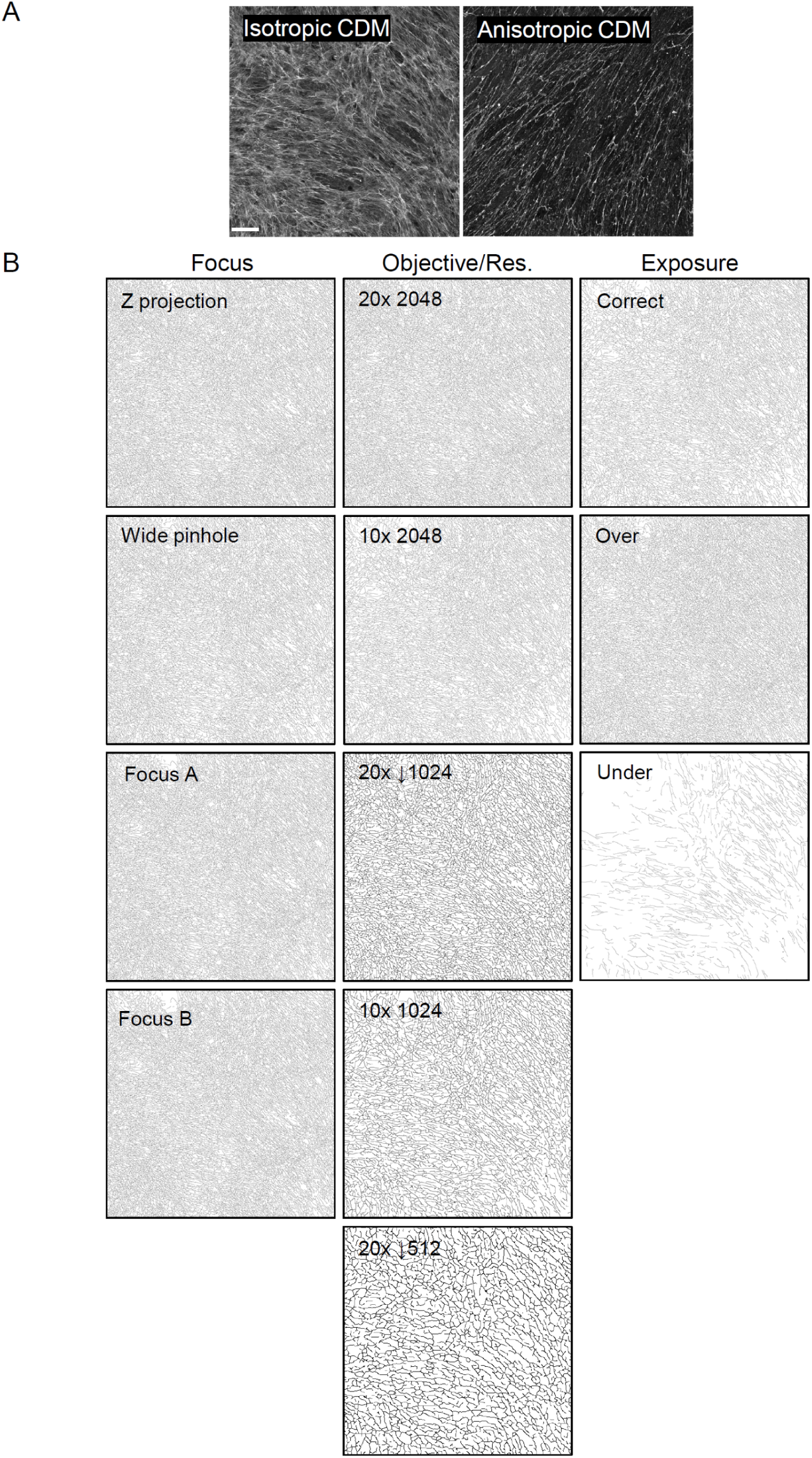
(A) Examples of the images of the isotropic and anisotropic matrices used for robustness testing. Images are a single confocal section captured with a 20x 0.75 NA objective containing 2048 × 2048 pixels, corresponding to 850 microns x 850 microns. Scale bar is 100 microns. (B) Masks are shown for the filament network generated from images captured with varying focal plane, objective (10x or 20x), pixel sizes, and exposure. Downward arrow indicates that the image was downsized to the indicated number of pixels after acquisition, but prior to generating the mask.

**Supplementary Figure 2.**
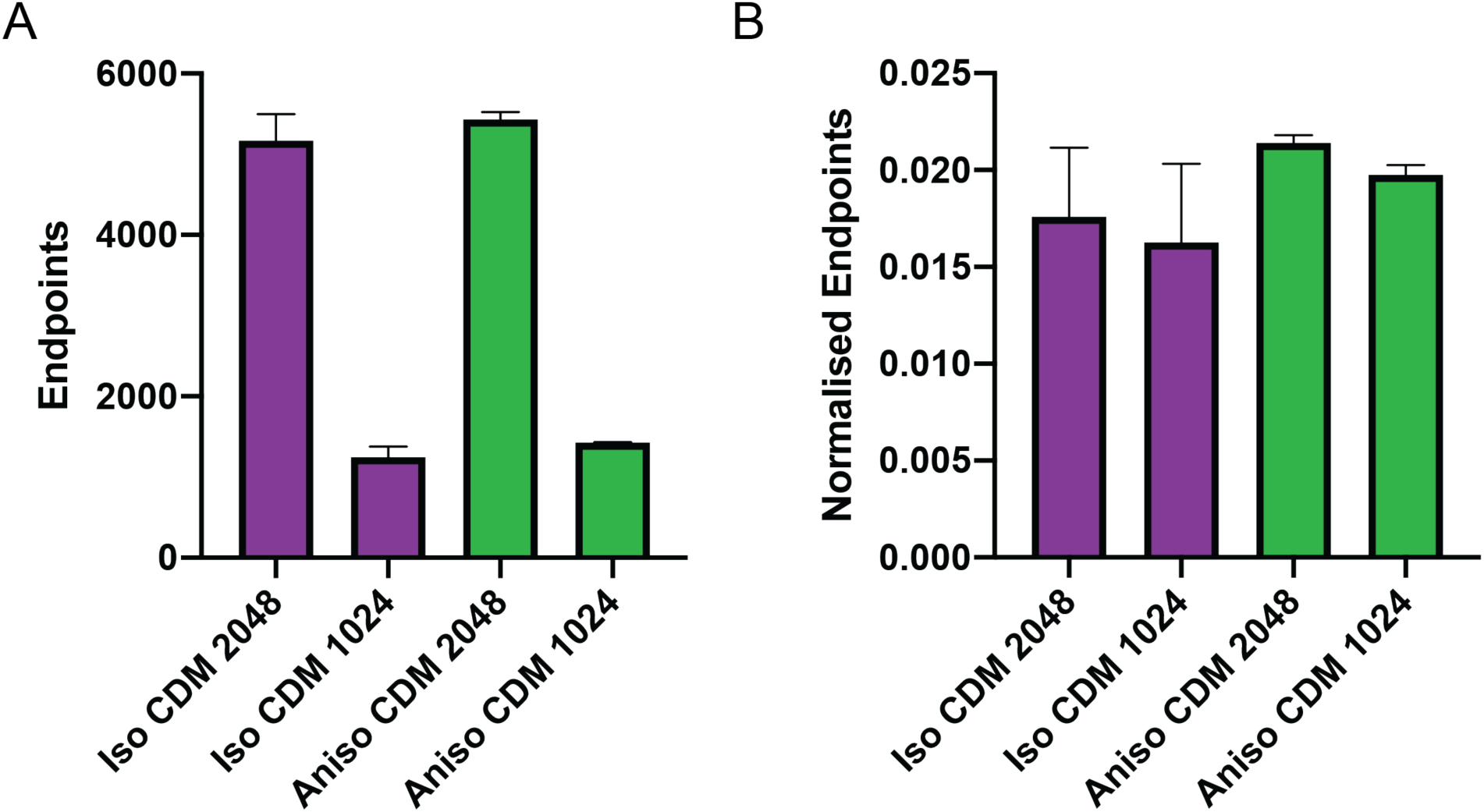
Effect of normalisation. (A) Shows quantification of endpoints from two images with 2048 pixels spanning 850 microns and two images with 1024 pixels spanning the same distance. Purple indicates Isotropic CDM and green indicates Anisotropic CDM. (B) Shows quantification of endpoints/length from two images with 2048 pixels spanning 850 microns and two images with 1024 pixels spanning the same distance. Purple indicates Isotropic CDM and green indicates Anisotropic CDM.

**Supplementary Figure 3:**
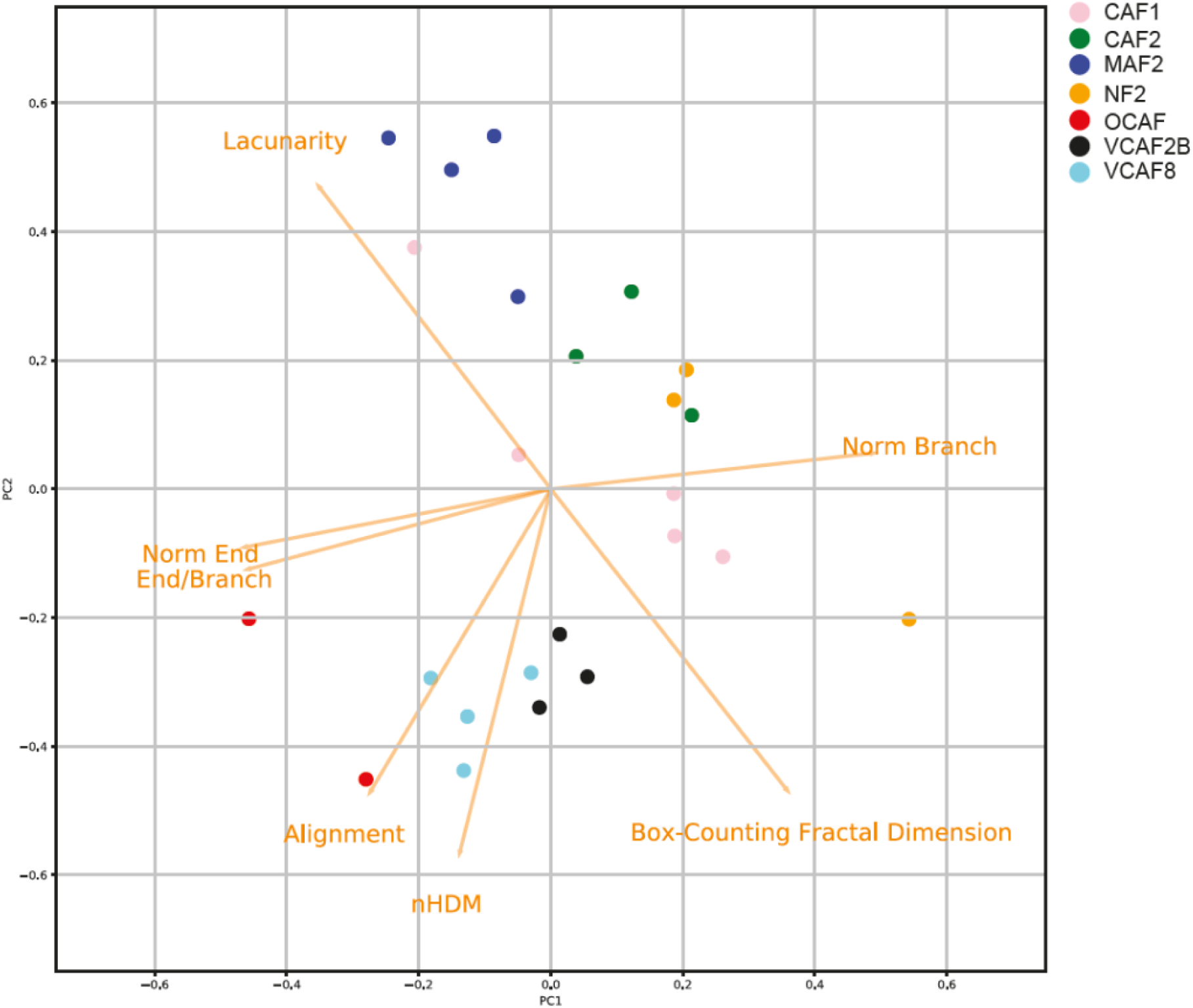
Discrimination of different cell-derived matrices. Image shows a PCA plot of the metrics of CDMs generated by seven different fibroblasts.

**Supplementary Figure 4.**
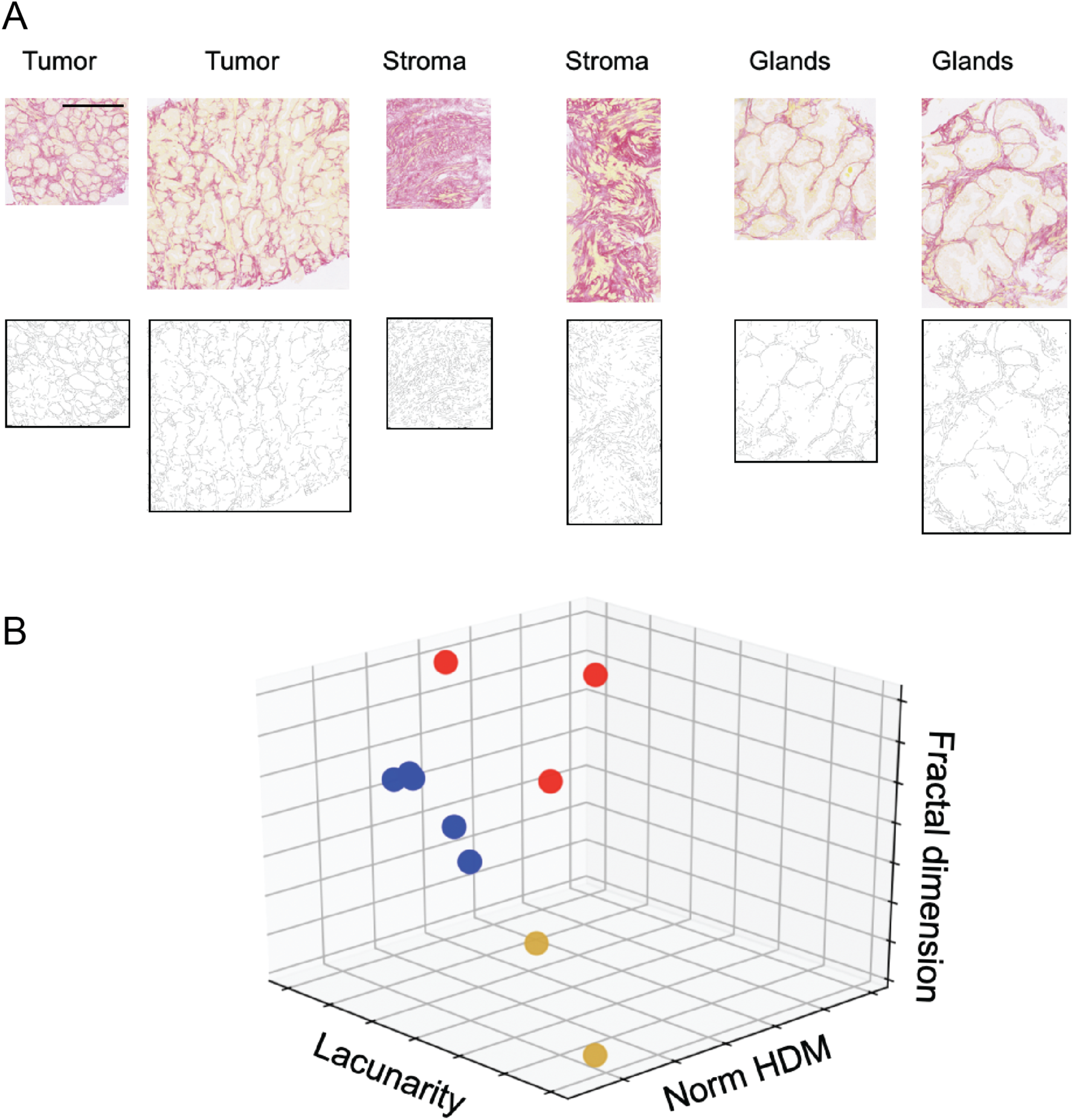
Quantifying prostate cancer images. (A) Normal prostate and tumour biopsies stained with picrosirius red are shown above their corresponding mask. Scale bar is 250 microns. Normal prostate images are sub-divided into glandular or stromal regions. (B) Three-dimensional plot showing the separation of normal glands (yellow), normal stroma (red), and tumour (blue) based on normalised HDM, fractal dimension, and lacunarity metrics.

**Supplementary Figure 5:**
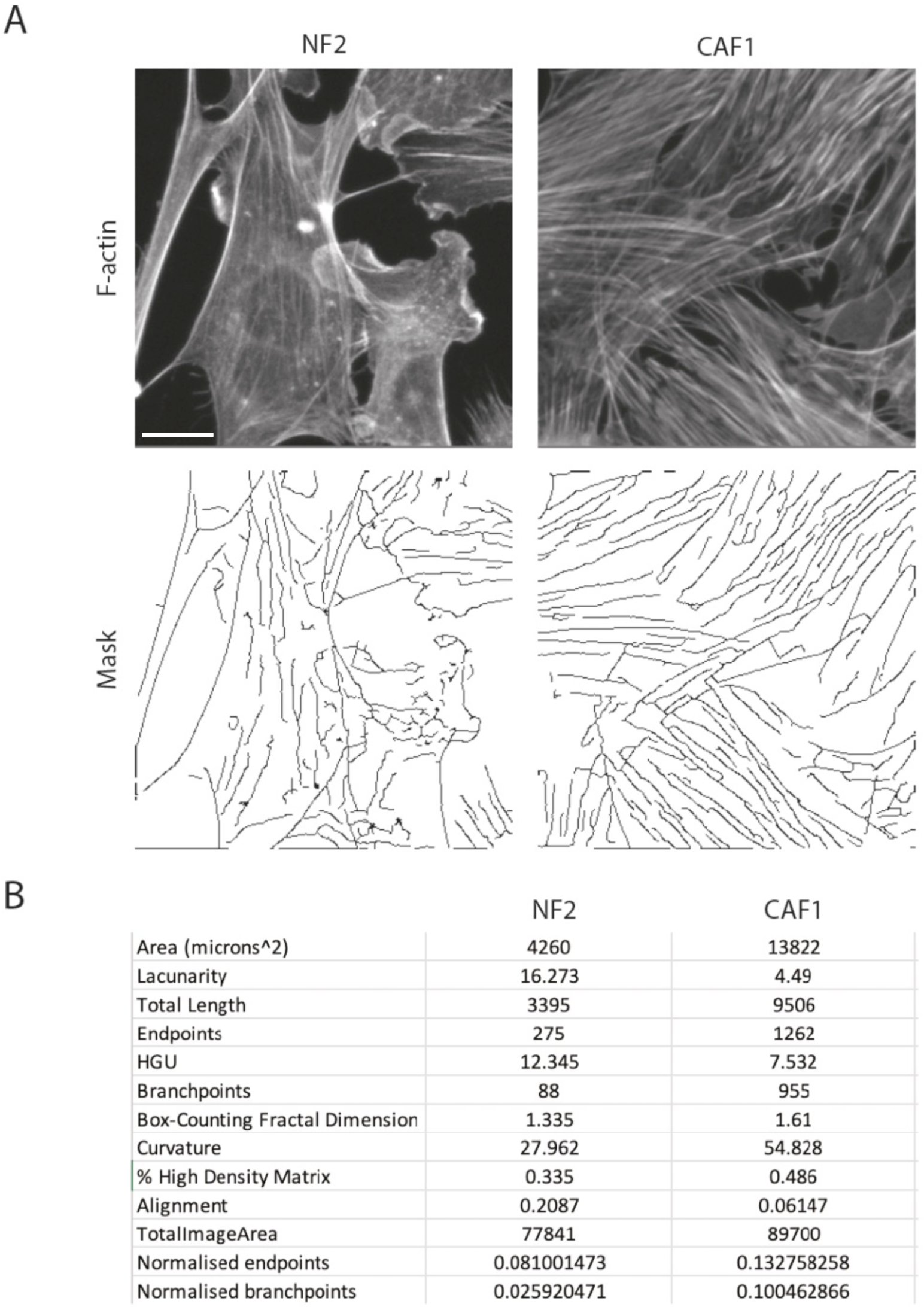
Applications of TWOMBLI to actin cytoskeleton. Images show the F-actin cytoskeleton from NF2.1 (fourth images from left) and CAF1 (left-hand panel) growing on 12kPa fibronectin coated substrate (images adapted from Calvo et al Cell Reports 2015 ^31^). Scale bar is 10 microns.

